# Functional impact of the hyperduplication genomophenotype in high copy number endometrial cancer

**DOI:** 10.1101/2024.09.20.613693

**Authors:** Angela Florio, James Smadbeck, Sarah H. Johnson, Wan-Hsin Lin, Dorsay Sadeghian, Sotiris Sotiriou, Rebeca Salvatori, Ryan W. Feathers, Taylor Berry, Lindsey Kinsella, Faye R. Harris, Alexa F. McCune, Stephen J. Murphy, Mohamed F. Ali, Abdulmohammad Pezeshki, Michael T. Barrett, Leah Grcevich, Ilaria Capasso, Luigi Antonio De Vitis, Gabriella Schivardi, Tommaso Occhiali, Alyssa M. Larish, John Weroha, Mitesh J. Borad, John Cheville, Panos Z. Anastasiadis, Andrea Mariani, George Vasmatzis

## Abstract

High copy number endometrial cancers (HCNEC) are dominated by excessive duplications scattered across the genome, termed here as the **H**yper**D**uplication **G**enomo**P**henotype (HDGP). Although correlated with cancer progression, its biological significance and implications for therapy have not yet been established. We identified locations and sizes of duplications in 171 endometrial cancer cases and designated 71 HCNEC cases as HDGP. We also investigated the response to the pan-ERBB inhibitor afatinib in a subset of HDGP-EC cases with *ERBB2/ERBB3* duplications using a patient-derived three-dimensional culture model. Our analysis demonstrates that beyond tandem duplications there is a more general pattern involving coordinated duplication of multiple distant regions of the genome, demonstrating preferential selectivity to over-expressed potential oncogenes within a broad network. This suggests that HDGP increases tumor fitness and resistance to therapy by perturbing important gene networks in concert rather than only driver genes, suggesting a mechanistic basis for the ineffectiveness of targeted drugs in these patients and highlighting the need for combination therapies in these highly aggressive cases.

## Introduction

Historically, roughly 80 percent of endometrial cancer (EC) cases were classified phenotypically as type I while approximately 20% were classified as type II^(1)^. Type I ECs are considerably less aggressive than type II and often can be successfully treated via hysterectomy alone; so while type I ECs account for a much larger percentage of cases, type II ECs account for significantly more deaths due to recurrence of disease (∼50%)^(2)^.

More recently, endometrial cancers have also been subtyped via genomic alterations into four classifications: POLE ultramutated, microsatellite instability hypermutated, low copy number, and high copy number^(3)^. While the first three subtypes are predominantly type I, the high copy number subtype are typically type II. It is also known that the high copy number subtype has a high incidence of what has been called a Tandem Duplication Phenotype (TDP) and has also been observed in ovarian, gastric, and breast cancer ^(4,5)^. The TDP has been described extensively by Menghi and Liu from the Jackson Laboratories ^(4,6,7)^, classifying TDP tumors by tandem duplication size into three classes: small (∼11 kb), medium (∼231 kb) and large (∼1.7 Mb).

Whole-genome sequencing (WGS) technology can identify both structural variants (SVs) and copy number variants (CNVs). Structural variants are large chromosomal rearrangements such as tandem duplications, deletions, inversions, and chromosomal translocations that result from the aberrant joining of disparate regions of the genome (DNA junctions). Mate-pair sequencing (MPseq) and paired-end sequencing are designed to determine copy number variants and the junctions resulting from these structural variants with high accuracy and resolution, and allow observation of complex events at the whole-genome level ^(8–11)^.

To better understand the pattern and impact of duplications in endometrial cancer, we analyzed MPseq and WGS data from 171 EC samples. This group is biased toward high-risk cases because many of them were collected through an initiative that focused on aggressive cancers. Through analysis of the structural variants found in each case we were able to identify a pattern of recurrent tandem and non-tandem duplications we termed the Hyper-Duplication Genomophenotype (HDGP). The patterns of distribution of the duplications across the genome revealed their oncogenic potential and allowed the construction of a network of genes implicated in endometrial cancer progression.

## Results

### The hyper-duplication genomophenotype in EC contains both tandem and concatenated duplications

Structural variant analysis (SVA) ^(8)^ and copy number variant (CNV) analysis ^(9)^ of the available EC data set were used to detect structural genomic abnormalities in EC cases. SVA revealed that about one third of the cases contained numerous aberrant DNA junctions due to rearrangements. An example is shown in Figure 1, where magenta lines represent the junctional rearrangements in this tumor’s genome. CNV analysis of this case detected several large deletions such as the 3p arm, and gains such as the 4p arm (Supplemental Fig. S1).

**Figure 1.**
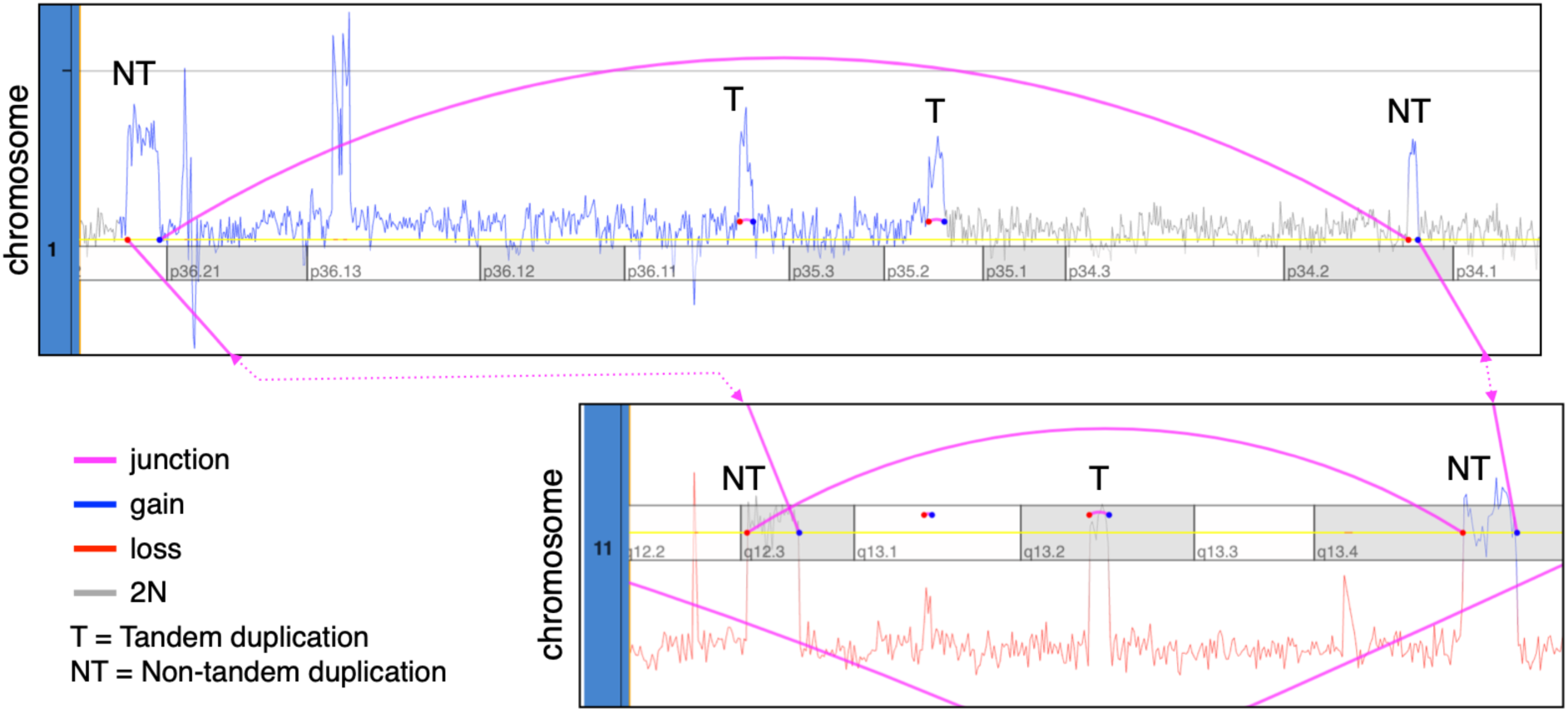
A hyperduplicated EC case with both tandem and non-tandem duplications. A tandem duplication will have a short elevation due to a CNV gain and a rearrangement line that signifies the junction. Tandem duplications (T) appear as a small copy number peak (blue) formed by adjacent duplications linked by a short junction (magenta), while non-tandem duplications (NT) are non- adjacent peaks linked by a longer junction. Tandem and non-tandem duplications occurring in 1N regions (red) bring the duplicated regions up to 2N (grey), although with a loss of heterozygosity. The extremities of the junction lines are colored according to the mapping direction (red if reverse, blue if forward). The dotted magenta lines show where the junctions connect chromosomes 1 and 11. A whole-genome view is available in Supplemental Fig. S1.

Additionally, CNV analysis detected many small duplications evident by elevated small regions across the genome. Most duplications were tandem as defined by the junctions spanning their edges (Fig. 1). Interestingly, there were a significant number of small duplications associated with junctions generated by complex rearrangements involving multiple distant locations of the genome. Following the junctions of these duplications reveals that small segments were duplicated and concatenated in sets of two to five, creating contiguous DNA fragments; the fragments are either integrated into one of the genomic locations involved or are associated with extra chromosomal elements. Structurally, these duplications resemble what have previously been described as “templated insertions” ^(12)^; however, templated insertions have been strongly attributed to replication in a BRCA2 deficient environment and would preclude the possibility of extrachromosomal elements ^(13)^. Given the absence of BRCA2 mutations in our data set, we cannot definitively classify our concatenated duplications as templated insertions. To identify hyper-duplicated cases for further analysis, we developed an algorithm (see Methods) that incorporate junction and read-depth information to separately identify tandem and concatenated duplications in 171 EC cases.

Menghi et.al. ^(7)^ grouped tandem duplications into small (∼11kb-class 1), medium (∼231kb- class 2), and large (∼1.7Mb-class 3). Our SNV pipeline calls intra-chromosomal events larger than 30kb due to the larger fragment size used by MPseq, so we focused on medium and large duplications at thresholds of 51 kb-6400 kb (classes 2 and 3). The distribution of duplication counts for all cases defined a minimum threshold of approximately 20 duplications for the hyper-duplication genomophenotype in EC (Supplemental Fig. S2A). All cases with the hyper-duplication genomophenotype contained both tandem and concatenated duplications, with strong ∼2.4:1 correlation (Supplemental Fig. S2D) as calculated by logistic regression.

### Recurrently duplicated genomic locations are not random

Menghi, et al. argued that the duplication events within the HDGP are not random but tend to hit genes of functional importance. To investigate whether the identified duplications occurred preferentially in any genomic locations in the EC cases, we segmented the genome into 25 kB consecutive windows, then overlaid 51kB-6400kB duplicated regions from all HDGP cases and used sample statistics to calculate the duplication frequency of each genomic location. To establish statistical significance, we ran 1,000 simulations by constructing sets of duplication events with lengths and sizes identical to those observed in the EC samples and distributing them randomly across the genome to determine the frequency of overlap. From the random simulations we determined that recurrent bins of 10 or more duplications are highly unlikely with the given set size, with a probability of less than 0.005 using area under curve. We then compared this simulated level of overlap to the level of overlap between duplications that was observed in our 71 HDGP-EC samples (Supplemental Fig. S2B). Interestingly, there were many regions that were duplicated 10-24 times in our data set. Thus, we conclude that the locations of duplications are not random, but rather demonstrate preferential selectivity for specific regions of the genome.

### Recurrently duplicated genomic locations of cases with the hyper-duplication genomophenotype are enriched in oncogenes

We investigated whether recurrent duplicated areas contain oncogenes with increased activity, either due to the duplication event or as the result of fusions resulting from junctions. To determine if duplications preferentially contain any cancer genes, we first combined OncoKB ^(14)^ and COSMIC ^(15)^ databases and classified genes as oncogenes, tumor suppressors, or undetermined based on each database’s criteria. Close analysis of the **R**ecurrent **Mi**nimally **D**uplicated **R**egions (**RMDRs**) across the genome revealed that 36 oncogenes including *ERBB2, ERBB3, MYC,* and *ESR1* were preferentially duplicated (Supplemental Fig. S2C, Supplemental Fig. S3). However, 14 tumor suppressors were also duplicated more frequently than expected (>9 times). Statistical analysis using the Wilcoxon rank sum test with continuity correction revealed significant oncogene preferential duplication compared to tumor suppressors (p = 2.1e-07). A closer look at tumor suppressors most frequently duplicated showed that they were located proximal to oncogenes that were also preferentially duplicated. This suggested that frequency of duplication alone should not be used to pinpoint the EC-related genes. Additionally, there were many RMDRs that were void of recognized oncogenes but contained genes with oncogenic potential based on their function and participation in EC-related pathways. We investigated all RMDRs with 10 or more recurrent duplications and selected genes that had evidence of over-expression in endometrial cancer and/or were implicated in the endometrial cancer literature (Supplemental Fig. S3).

### Recurrently duplicated genomic locations of cases with the hyper-duplication genomophenotype could point to potential EC oncogenes

A tier system was developed to allow deeper analysis through hierarchical grouping based on frequency of duplication and significance of expression compared to the non-EC data set. Each gene was evaluated individually in the context of its genomic region and was assigned a preliminary tier based on its duplication frequency in the HDGP-EC data set. Tier assignments were further refined up or down based on expression t-tests compared to non-EC cancer cases in the overall data set (Supplemental Fig. S4A). The tier groupings resembled a normal distribution, with Tier 2 and Tier 3 together comprising 64 percent of the overall list (Supplemental Fig. S4B,C). Overlapping distributions show how the median frequency of duplications decreases from 14 (Tier 1) to 10 (Tier 4), and that Tier 1 shows a bimodal distribution with a second, smaller population with frequencies around 18-20 (Supplemental Fig. S4E). The distribution of certain chromosomes across the tiers was significantly different from that of the overall gene list (Supplemental Fig. S4D); chromosome 12 was the only distribution skewed toward Tiers 1 and 2, likely due to a broad peak extending from 12:55954104 to 12:56224341 that includes *CDK2, ERBB3, PA2G4*, and neighboring genes in addition to five separate peaks.

Genes from each tier were functionally annotated for Gene Ontology: Biological Process, and annotations were categorized into broader groups. The distribution of these broader groups across tiers was evaluated against the overall distribution of genes to determine which processes and/or functions were overrepresented in a particular tier (Fig. 2A). Apoptosis, Proliferation, and Migration were significantly overrepresented in Tier 1, supporting our finding that oncogenes are preferentially duplicated. Signal Transduction was overrepresented in Tiers 2 and 3, while the distributions of Transcription and Metabolism were not significantly skewed.

**Figure 2.**
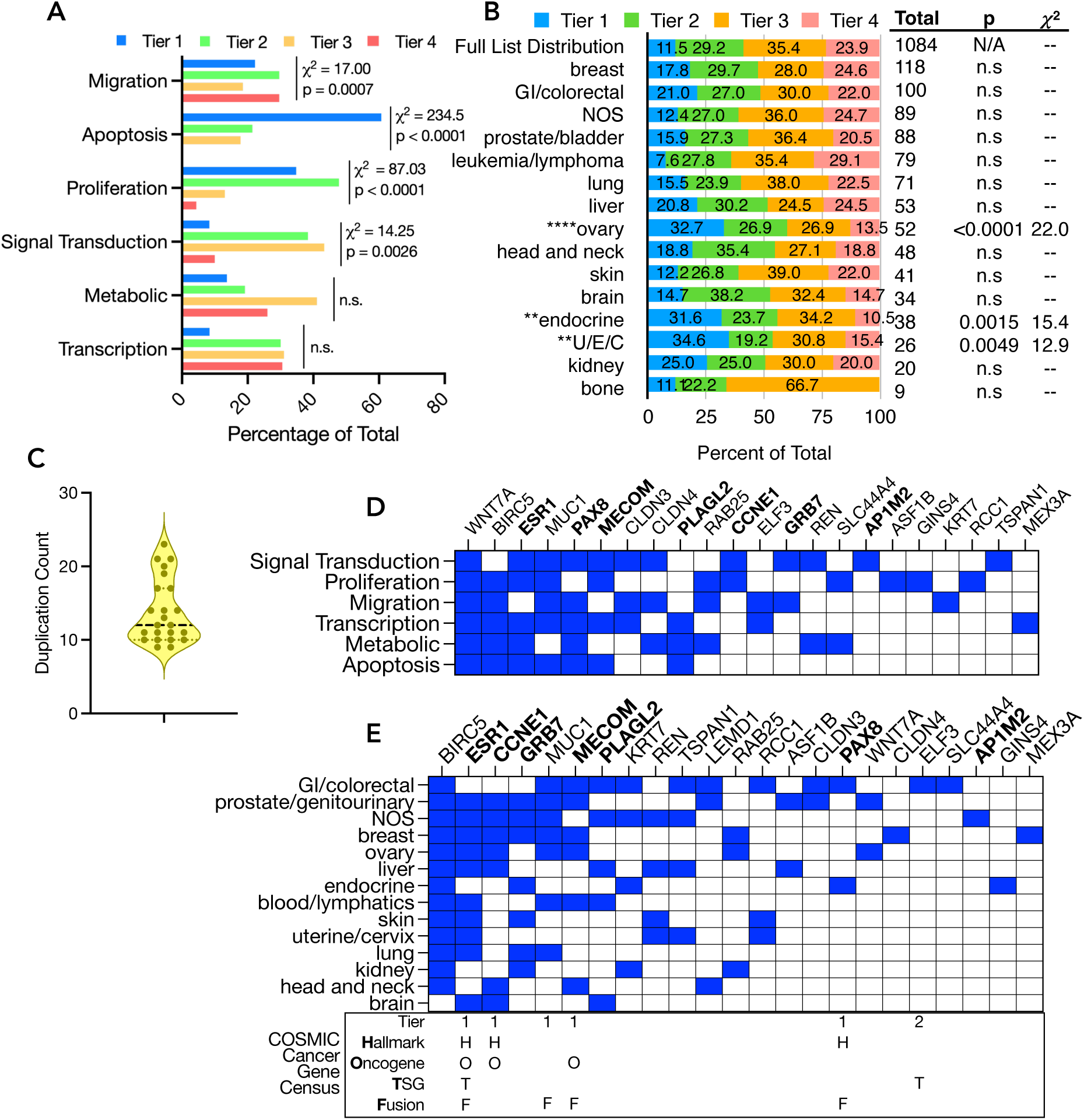
Functional and cancer-specific annotation of hyperduplicated genes. **A)** Hyper-duplicated genes were functionally annotated using Gene Ontology: Biological Process, and annotations were sorted into broader functional groups. Chi-square test, alpha = 0.05, DF = 3. **B)** Each tier was annotated with cancer-specific associations from DisGeNET and GAD Disease, and the annotations were sorted into broader groups based on tissue/system. U/E/C represents uterine, endometrial, and cervical cancers. Chi square test, alpha = 0.05, DF = 3. **C)** The Top 23 genes show variation in duplication count, reflecting the sorting criteria. **D)** The Top 23 genes were functionally annotated using Gene Ontology: Biological Process, and annotations were sorted into broader functional groups. **E)** The Top 23 genes were annotated for tissue-specific cancer associations using DisGeNET, GAD Disease, FABRIC, and the Atlas of Genetics and Cytogenetics in Oncology and Haematology. ESR1, CCNE1, MUC1, MECOM, PAX8, and ELF3 are included in the COSMIC Cancer Gene Census. Our Top 7 Genes are in bold.

Cancer associations for the genes of each tier were compiled using DisGeNET ^(16)^ and the Genetic Association Database ^(17)^, and the annotations were broadly grouped by tissue. The distribution of tiers across tissue-specific cancer associations was investigated to identify possible similarities to other cancers (Fig. 2B). Tier 1 had the highest number of associations overall, and was significantly overrepresented in association with ovarian, endocrine, and uterine/endometrial/cervical cancers.

### Integrated pathway analysis suggests the involvement of multiple linked pathways and a transcriptionally active environment supporting proliferation and invasion

Tier 1 was re-evaluated separately with more stringency to identify the most statistically important genes: those appearing in at least 15 cases, with expression statistics probability and t-statistic less than 10-10. Seven genes met these criteria: transcription factors *ESR1*, *MECOM*, *PAX8*, and *PLAGL2*; adapter proteins *AP1M2* and *GRB7*, and cyclin *CCNE1* (Table 1). *ESR1* was duplicated as the sole peak in region 21 (6q25); *MECOM* was duplicated as part of region 15 (3q26), which also contains *PRKCI*; *PAX8* was duplicated as part of region 11 (2q14), which includes a cluster of interleukins and interleukin receptors; and *PLAGL2* was duplicated as part of region 58 (20q11), which includes *BDL2L1*, *E2F1*, *DNMT3B*, *TPX2*, *ID1*, *MYLK2*, and *TTLL9*. *GRB7* was duplicated as part of region 49 (17q12), which also includes *ERBB2*, *MIEN1*, *SNORA21*, *LINC00672* and junctional protein *JUP*; *AP1M2* was duplicated as part of region 54 (19p13), which includes *CARM1*, *CDKN2D* and many other genes with high frequency but low significance, suggesting known cancer-related genes; and *CCNE1* was duplicated as part of region 57 (19q12), which includes ovarian cancer-associated genes *POP4*, *URI1*, *C19orf12*, and *PLEKHF1* ^(16)^ (Supplemental Fig. S5).

**Table 1.**
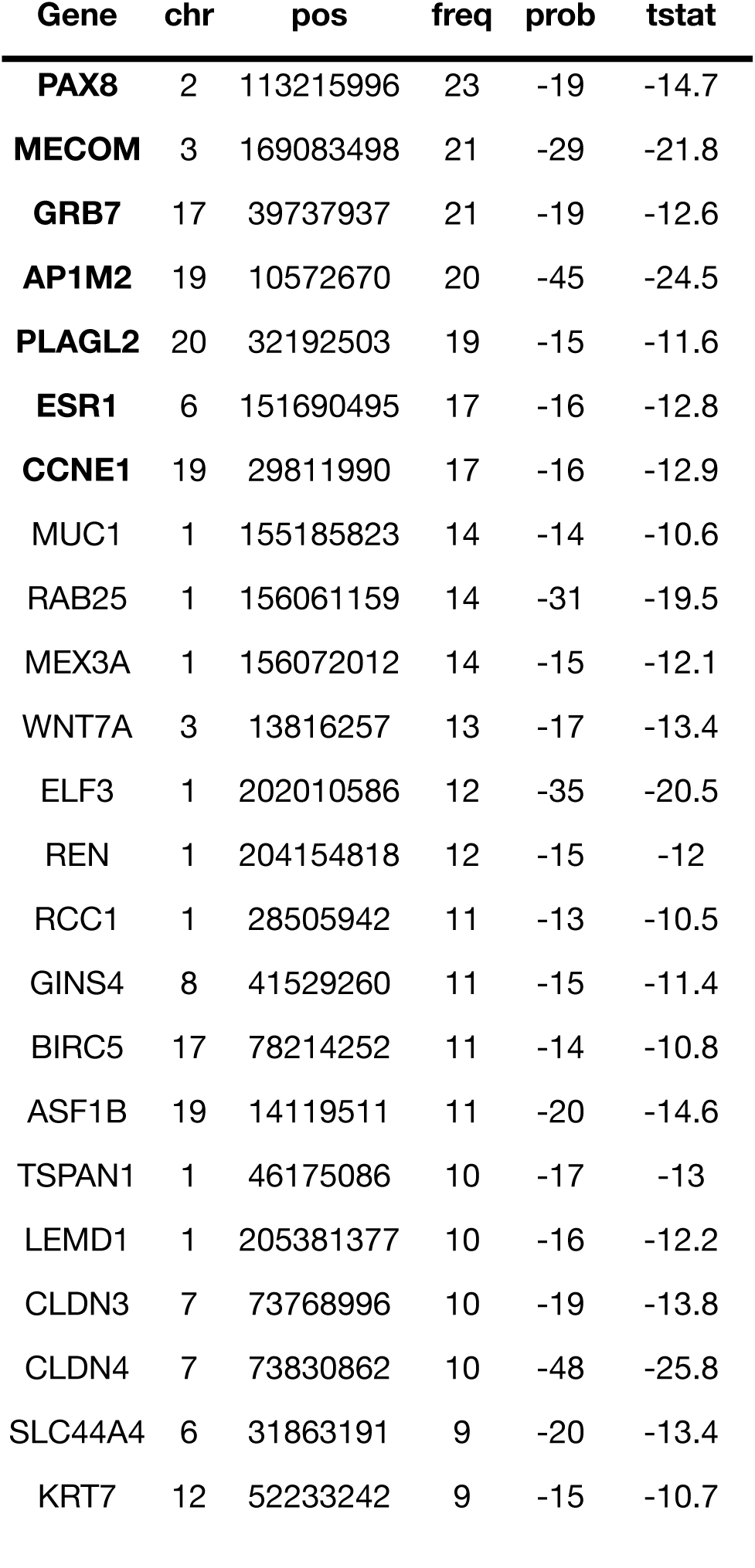
The Top 23 genes include the Top 7 Genes (bold), which are duplicated at the highest frequency and have the most significant expression statistics in our HCNEC data set, as well as an additional 16 genes that met the more stringent expression criteria but appear at lower frequency. chr indicates chromosome; pos indicates position on the chromosome; prob and tstat indicate expression statistics.

An additional 16 genes were identified as meeting the expression statistic requirements but appearing in fewer cases (Table 1). Together, these Top 23 Genes were re-evaluated as a set. The genes were functionally annotated as before and categorized into broader groups (Fig. 2D). *WNT7A* was the only gene with annotations in all six groups; *BIRC5*, *ESR1*, *MUC1*, and *PAX8* had annotations in five groups. This suggests that these genes exert the broadest influence over cell behavior, although this does not imply any magnitude of impact as the effect in any or all groups may be minimal. The Top 23 Genes were annotated for tissue-specific cancer associations as before and combined with additional data from FABRIC ^(18)^ and the Atlas of Genetics and Cytogenetics in Oncology and Haematology ^(19)^. As a group, these genes were most highly associated with gastrointestinal and colorectal cancers. Interestingly, only five genes were associated with cancers of the uterus or cervix. Only 6 of our Top 23 Genes – *ESR1*, *CCNE1*, *MUC1*, *MECOM*, *PAX8*, and *ELF3* – are included in the Catalog of Somatic Mutations in Cancer (COSMIC) Cancer Gene Census ^(15)^ (Fig. 2E).

The gene list was mapped to an integrated pathway representing cancer cells’ major routes to proliferation, survival, and invasion (Fig. 3); redundancies were removed to preserve readability. Focal adhesion kinase (*PTK2*) and Akt (*AKT1*) emerged as hubs; focal adhesion kinase transduces signals from cell surface receptor tyrosine kinases (RTKs) and cell-matrix interactions to effect actin cytoskeleton reorganization and invasion signaling, while Akt receives signals from RTKs, Notch, and Wnt via PI3K to effect cell proliferation and survival (Fig. 3A). It is worth noting that although the frequency of *AKT1* duplications fell just below our significance cutoff, it was included here to illustrate the flow of information because it is known to be important in cancer signaling even if it is not duplicated.

**Figure 3.**
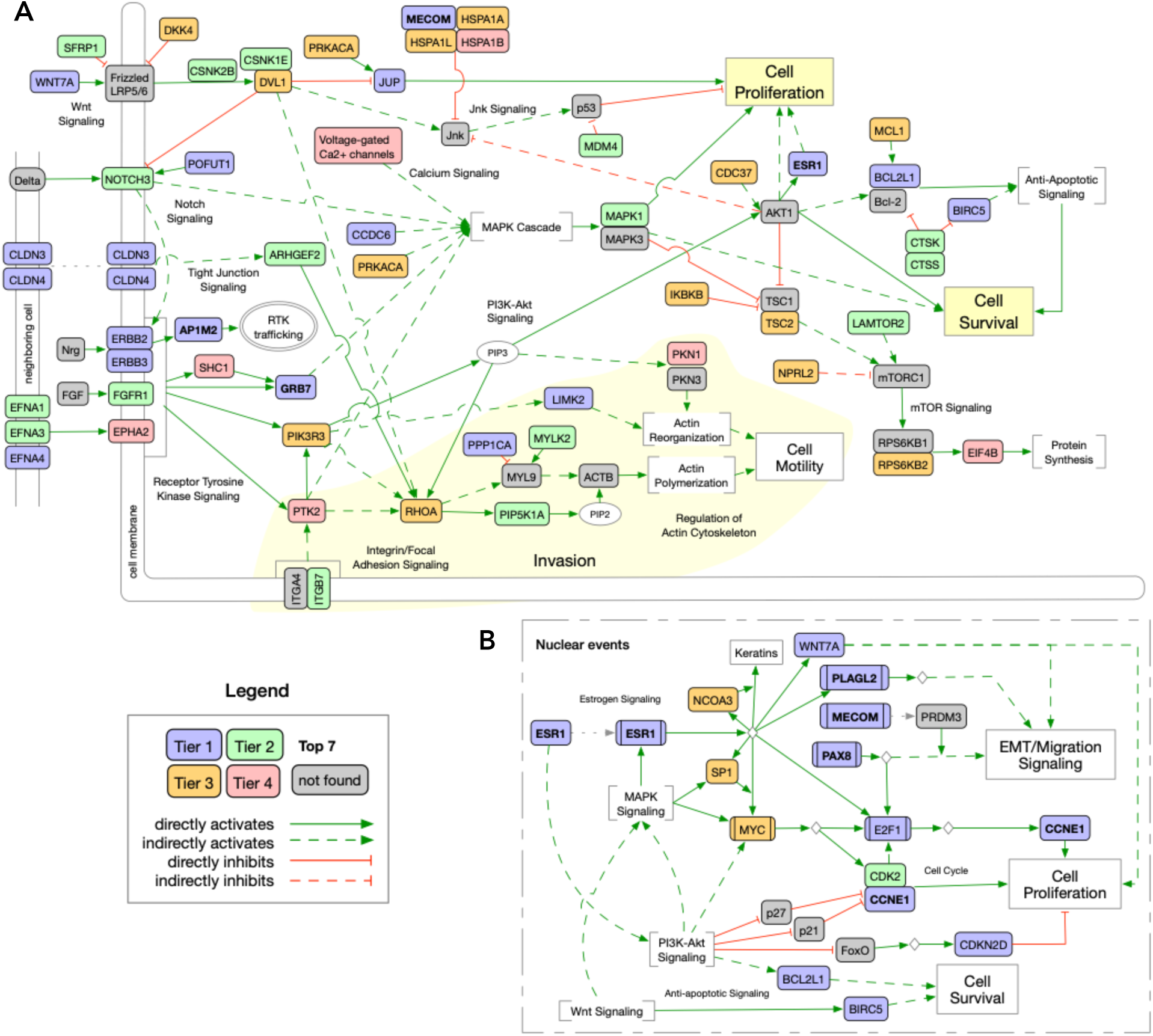
Hyperduplicated genes were mapped to common cancer pathways. **A)** Signaling through receptor tyrosine kinases (RTK), Notch, and Wnt induces proliferation and apoptosis avoidance through PI3K-Akt and MAP kinase pathways. Invasion-related signaling is represented through integrin activation of focal adhesion kinase (*PTK2*) and downstream through RhoA. Focal adhesion kinase signaling indirectly reinforces signaling through the MAPK and PI3K-Akt pathways. Motility signaling is reinforced by Notch, Wnt, PI3K-Akt, and Tight Junction signaling through activation of RhoA as well as indirect PIP3 activation of actin reorganization. **B)** *ESR1*-induced transcription appears to dominate nuclear events. Supported by incoming signal from *MAPK*, *PI3K-Akt*, and *Wnt* signaling, this results in transcription of genes involved in cell migration, proliferation, and survival. Redundancies have been removed and localization is approximate to preserve readability. Nodes are colored by tier; Top 7 genes are in bold; genes in gray were not significantly duplicated, but were included for context. Green edges indicate activation, red edges indicate inhibition; solid edges indicate direct activity, dashed edges indicate indirect activity.

Estrogen receptor (ESR1)-mediated transcription appears to dominate the nuclear events (Fig. 3B), initiating transcription of its own cofactors *NCOA3* and *SP1*, which in turn initiates transcription of *MYC*. Myc (*MYC*) initiates transcription of *E2F1*, *CDC25A*, and *CDK2*, leading to transcription of *CCNE1* and continuation of the cell cycle, while signaling through the PI3K-Akt and Wnt pathways support anti-apoptotic signaling through BCL2L1 and BIRC5, respectively. PI3K-Akt signaling removes repression of CDK2/CCNE1 and inhibits the transcription of *CDKN2D*, supporting cell proliferation. Pax-8 (*PAX8*) reinforces proliferative transcription by inducing transcription of *E2F1* and participates in migration-promoting transcription with *PLAGL2*. Cell migration and epithelial-mesenchymal transition (EMT) is stimulated by transcription products arising from *MECOM* isoform *PRDM3* acting as a coactivator for PAX8, as well as PLAGL2 targets via Wnt/beta-catenin signaling.

### Tandem duplication-associated ERBB2/3 endometrial tumors exhibit increased sensitivity to pan-ERBB inhibition in three-dimensional *ex vivo* culture

We next investigated the functional relevance of HDGPs by focusing on tumors harboring *ERBB2/3* duplications. Tumor cells harboring *ERBB2/3* tandem duplications exhibit increased expression in cases with *ERBB* duplications compared to cases without *ERBB2/3* duplication (Fig. 4A,B). Both expression of *ERBB2* (assessed by RNAseq) and expression of its protein product Her2 (assessed by immunostain) were higher in the *ERBB2*-duplicated cases when compared to the others (Fig. 4C). Using our previously established method ^(20,21)^, three-dimensional cultures of patient-derived tumor cells (microcancers) were generated from endometrial tumors and treated with afatinib for six days in a dose-response study. Afatinib is an irreversible pan-ErbB inhibitor that is approved for *EGFR*-altered non-small cell lung cancer (NSCLC), and its anti-neoplastic effect in endometrial cancer with *ERBB2* alterations is actively investigated in both the pre-clinical and clinical settings ^(22,23)^. A shift in afatinib dose-response curves suggested that endometrial microcancers containing *ERBB2*/*3* duplications are more sensitive to afatinib, with significantly lower absolute half maximal inhibitory concentrations (IC50s) compared to microcancers with no *ERBB2/3* duplications (Fig. 4D,E). However, the *ex vivo* efficacy of afatinib at its reported Cmax of 0.052 mM was low, even in cases with *ERBB2/3* duplications, consistent with the existence of additional oncogenic drivers in these tumors.

**Figure 4.**
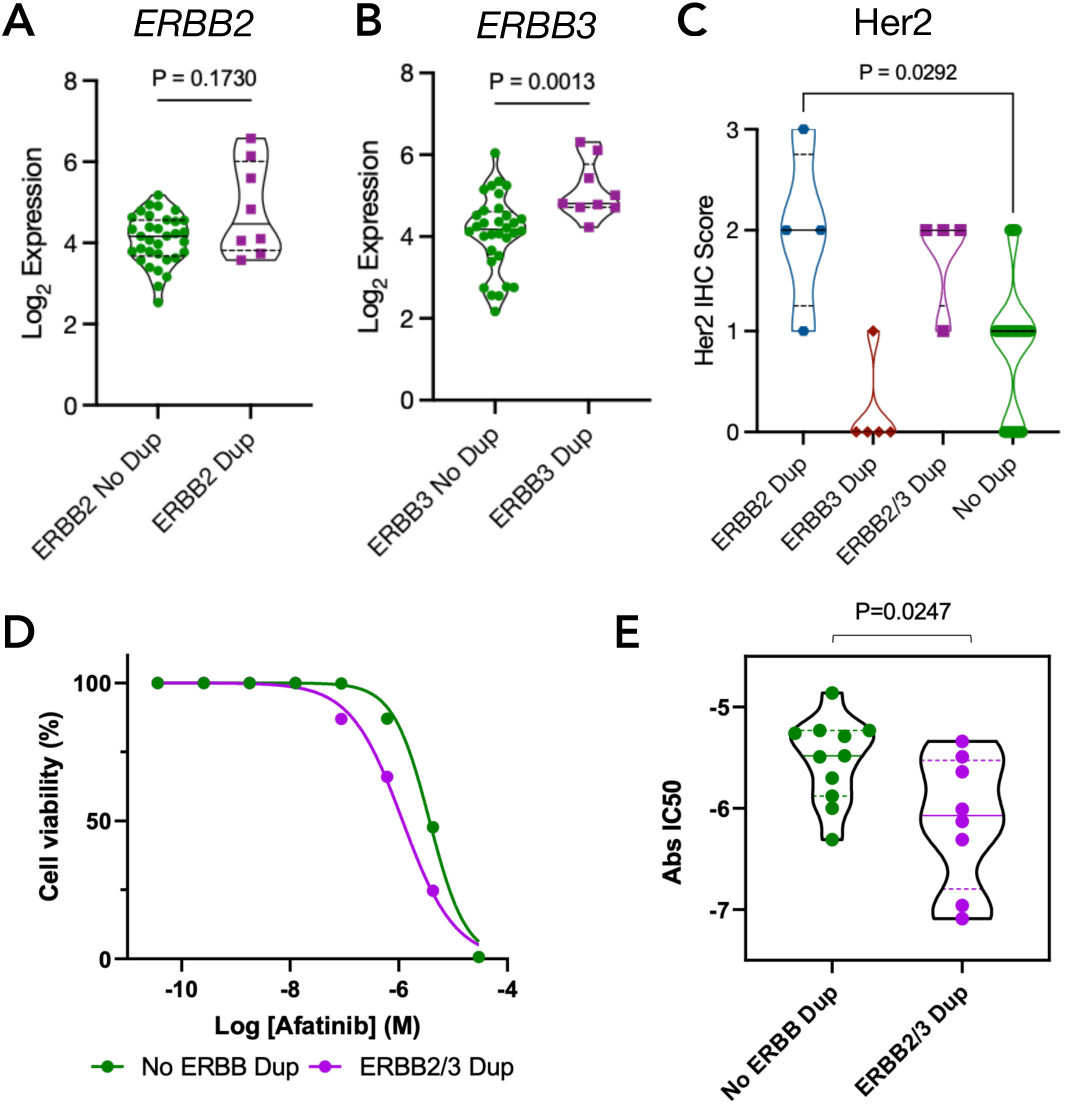
TD-associated ERBB2/3 endometrial microcancers exhibit increased sensitivity to pan-ERBB inhibition. **A, B.)** Truncated violin plots showing expression of ERBB2 (A) or ERBB3 (B) in cases with (purple) and without (green) ERBB2/3 duplications. Median (solid line) with quartiles (dashed lines), Mann-Whitney test. **C)** Truncated violin plot showing Her2 IHC scores for cases with duplications of ERBB2 (ERBB2 Dup, *n*=4), ERBB3 (ERBB3 Dup, *n*=5), both ERBB2 and ERBB3 (ERBB2/3 Dup, *n*=4), or neither (No Dup, *n*=23). Kruskal Wallis test with Dunn’s correction for multiple comparisons. **D)** Afatinib dose-response curves of endometrial microcancers. Microcancers were generated from patient tumors with (purple) or without (green) the presence of *ERBB2/3* duplication (Dup). Data points represent the median of data points from tested cases (no *ERBB2/3* Dup, *n*=11 cases; *ERBB2/3* Dup, *n*=5 cases). **E)** Truncated violin plot depicting absolute IC50 (Abs IC50) of afatinib in endometrial microcancers with (purple) or without (green) the presence of *ERBB2/3* duplication (no *ERBB2/3* Dup, *n*=11 cases; *ERBB2/3* Dup, *n*=8 cases). Median (solid line) and quartiles (dashed line) are indicated.

## Discussion

Here we present a characterization of the duplication variants identified in high copy number endometrial cancer by MPseq and paired-end WGS. We demonstrated that what has been termed the TDP, is in fact a more general duplicator phenotype that includes distant intra- and inter-chromosomal duplications events often involving 2-5 distant genomic locations. This suggests that WGS analysis of samples for the duplicator phenotype would risk undercounting the levels and locations of duplications in a sample if tandem duplications alone were isolated for analysis. Instead, our results suggest that the criteria for inclusion in analysis should be expanded to include any region of focal gain that is supported by a DNA junction regardless of strand orientation. Unfortunately, from the junction data and the shallow sequencing it is impossible to perform a de novo assembly to determine if the non-tandem duplicated regions are integrated in the chromosomes or are extrachromosomal.

Through an analysis of the recurrent duplications found in HCNEC samples we concluded that the duplications were not randomly distributed across the genome. Simulations of the random distribution of duplications of similar size demonstrated that the recurrent overlap observed in our samples deviated significantly from expected. Recurrent duplications across samples contained genes associated with important cancer pathways. This suggests that the accumulation of duplications may not be random or simply a sign of an unstable genome, but rather a perturbation of important genetic pathways that promotes tumor progression, enhances tumor fitness, and increases therapeutic resistance and aggressiveness of a cancer; this is supported by the fact that the particular tumor types that present with this phenotype (i.e.; serous carcinomas and triple negative breast cancers) are known to behave aggressively and be resistant to most chemotherapies. Such a mechanism also presents difficulties surrounding targeted treatment of a cancer that evolves by activating multiple tumorigenic pathways through widespread duplication of genes. This is corroborated by the response of *ERBB2/3* duplicated tumors to afatinib treatment *ex vivo*. A shift in dose-response curves and a reduction in absolute IC50 values for afatinib monotherapy argues that the increased expression of *ERBB2/3* as a result of hyper-duplication contributes to tumor growth. However, the underwhelming response to afatinib at its reported Cmax indicates the presence of multiple oncogenic drivers in HDGP- ECs.

As with any cancer, endometrial cancer is a disease characterized by the accumulation of genetic mutations over time. These genetic mutations result in uncontrolled growth due to disruption of normal cellular function. It is possible that the difference in outcome between the disease subtypes is a direct result of the differences between their genetic variations. Our analysis allowed for the construction of a signaling network for endometrial cancer. Several pathways were recurrently hit by the hyper-duplication phenotype; the ErbB and ER pathways were already known, and the cell cycle pathway was implicated via duplications of *CCNE1*, *CDK2* and *E2F* genes. Here we also introduce potential involvement of anti-apoptotic signaling via duplications of *BCL2L1* and *BIRC5*, as well as components of Notch signaling, Wnt signaling, RhoA signaling, and others.

We divided the gene list into tiers to allow us to identify the genes with the highest frequencies in our HCNEC dataset and the most significant expression differences with our larger, non-endometrial cancer data set. As the genes with the highest frequencies and the most highly significant expression levels compared to non-EC cases, *PAX8*, *MECOM*, *ESR1*, *GRB7*, *AP1M2*, *PLAGL2*, and *CCNE1* were named our Top 7 genes. *PAX8*, *MECOM*, *ESR1*, and *PLAGL2* are transcription factors with known cancer associations. Although canonically a thyroid-specific transcription factor, Pax-8 (*PAX8*) is a marker of serous endometrial carcinoma and has been shown to regulate cell growth through direct activation of *E2F1* expression ^(24)^. *MECOM* is an oncogene mostly associated with leukemias and lymphomas, although there is evidence of *MECOM* product *PRDM3* acting as a cofactor for Pax-8 transcription to promote expression of migration genes and sustain growth in ovarian cancer ^(25)^. *ESR1* encodes estrogen receptor alpha, a biomarker for breast cancer and endometrioid adenocarcinoma, although expression in serous endometrial cancer has been found to be less likely ^(26,27)^. *PLAGL2* is a weakly activating transcription factor that nonetheless has been found to induce epithelial-mesenchymal transition via Wnt/beta-catenin signaling in colorectal cancer and has been found to be over-expressed in many cancers ^(28,29)^.

*CCNE1* encodes cyclin E1, which controls the cell cycle G1/S transition by regulating Cdk2 (*CDK2*, Tier 2) activity. High levels of CCNE1 may lead to chromosome instability and contribute to tumor progression ^(30)^, and is associated with late-stage cancer and poorer prognosis ^(31,32)^. Although over-expression is common in a variety of malignancies, *CCNE1* has also been found to be amplified in up to 50% of high-grade serous ovarian and endometrial cancers ^(31,33–35)^, and has recently been shown to contribute to tumor cell proliferation and invasion in endometrial cancer ^(36)^.

*GRB7* and *AP1M2* encode adaptor proteins that function in receptor tyrosine kinase signaling and internalization. AP1-Mu2 (*AP1M2*) is a subunit of clathrin-associated adaptor protein complex 1, in which it mediates clathrin recruitment to the membrane and cargo molecule signaling recognition; it has also been shown to interact directly with ErbB proteins for internalization and trafficking ^(37)^. Grb7 is an adaptor protein shown to bind cell surface RTKs, including Her2 (*ERBB2*, Tier 1) and Her3 (*ERBB3*, Tier 1), to mediate downstream signaling ^(38,39)^; in this role, it also promotes cell migration by facilitating RTK signaling to focal adhesion kinase (*PTK2*, Tier 4) as well as downstream activation of MAPK and PI3K-Akt signaling. When not phosphorylated, Grb7 can bind to the 5’ UTR of target mRNA and repress translation similarly to miRNAs.

The duplicated region surrounding *GRB7* is interesting in that it includes several long intergenic non-coding RNAs (lincRNAs) and small nucleolar RNAs (snoRNAs) with known cancer associations. *SNORA21* (Tier 3) has been associated with distant metastases in gastric and colorectal cancers ^(40,41)^, and has been shown to promote proliferation in breast and colorectal cancer ^(41,42)^. *LINC00672* (Tier 4) has been previously reported to promote paclitaxel sensitivity in endometrial cancer by locally reinforcing p53 suppression of nearby gene *LASP1* (Tier 3), which contributes to tumor aggressiveness ^(43)^, and *LINC00974* has been found to interact with *KRT19* (Tier 3) to promote proliferation and metastasis through Notch signaling in hepatocellular carcinoma ^(44)^.

Many snoRNAs and lincRNAs remain functionally uncharacterized, but mounting evidence suggests complex roles in health and disease; our hyper-duplication regions contained several of each in addition to those mentioned above. snoRNAs are usually over-expressed in breast cancer, and have been shown to promote tumorigenesis ^(45)^; indeed, *SNORA44* (Tier 4) is a breast cancer oncogene ^(46)^, and *SNORNA21* (Tier 3), *SNORA73A* (Tier 3), and *SNORA73B* (Tier 3) have been shown to promote proliferation in breast and colorectal cancer ^(41,42)^. snoRNAs have also been found to repress tumor suppressor functions; *SNORNA80E* (Tier 2) has been shown to inhibit apoptosis through a p53-dependent pathway, and its expression levels are conversely correlated with p53 levels ^(47)^. Of the lincRNAs in our data set, *PVT1* (Tier 1) has been shown to co-amplify with *MYC* (Tier 3) ^(48)^ and *MALAT1* (Tier 2) is up-regulated in multiple cancers and its expression correlates with metastasis in early non-small cell lung cancer ^(3,49)^.

## Conclusions

Overall, the most important hyper-duplicated genes from a statistical perspective appear to promote an environment that supports proliferation, survival, and migration. This is to be expected, given that most of the duplicated genes are either known or potential EC oncogenes, but it also reinforces the idea that these duplications are advantageous and occur as part of the tumorigenic process rather than just random occurrences. Unfortunately, our network highlights the difficulty in treating HCNEC; there is no single driver to target, and there is strong representation of invasion-related signaling. While discussion of the effect of activating mutations such as *PIK3CA* and *TP53* fusion in EC is beyond the scope of this paper, future investigation may gain further insight by integrating RDMR analysis into a more comprehensive ‘omics study of HCNEC. Ultimately, further study of HDGP and associations with clinical outcomes could identify actionable targets and help better stratify this complex and heterogenous group of patients.

## Author Contributions

**A. Florio:** Conceptualization, data curation, formal analysis, investigation, methodology, project administration, software, validation, visualization, writing – original draft, writing – review & editing. **J. Smadbeck:** Conceptualization, data curation, formal analysis, investigation, methodology, software, validation, visualization, writing – original draft, writing – review & editing. **S. Johnson:** Formal analysis, methodology, software, validation visualization, writing – review & editing. **W.-H. Lin:** Data curation formal analysis, investigation, methodology, resources, software, validation, writing – review & editing. **D. Sadeghian:** Data curation formal analysis. **S. Sotiriou:** Data curation, formal analysis. **R. Salvatori:** Data curation, formal analysis, writing – review & editing. **R. Feathers:** Data curation, formal analysis, investigation, methodology, software, writing – review & editing. **T. Berry:** Investigation, writing – review & editing. **L. Kinsella:** Investigation, writing – review & editing. **F.R. Harris:** Investigation, methodology, writing – review & editing. **A. McCune:** Investigation, methodology. **S.J. Murphy:** Formal analysis. **M.F. Ali:** Investigation, methodology. **A. Pezeshki:** Writing – review & editing. **M.T. Barrett:** Formal analysis, resources, validation, writing – review & editing. **L. Grcevich:** Resources. **I. Capasso:** Resources. **L.A. DeVitis:** Resources, writing – review & editing. **G. Schiavardi:** Resources. **T. Occhiali:** Resources, writing – review & editing. **A.M. Larish:** Resources. **J. Weroha:** Data curation, resources, validation, writing – review & editing. **M.J. Borad:** Resources, validation, writing – review & editing. **J. Cheville:** Data curation, formal analysis, funding acquisition, resources, supervision, validation. **P.Z. Anastasiadis:** Formal analysis, funding acquisition, methodology, resources, software, supervision, validation, writing – review & editing. **A. Mariani:** Data curation, resources, supervision validation, writing – review & editing. **G. Vasmatzis:** Conceptualization, data curation, formal analysis, funding acquisition, investigation, methodology, project administration, resources, software, supervision, validation, visualization, writing – original draft, writing – review & editing.

## Acknowledgements

We would like to express our sincere gratitude to Giannoula Karagouga for her work with the endometrial cancer samples used to generate the data used in this project. This work was funded by Mayo Clinic Center for Individualized Medicine, Mayo Clinic Graduate School of Biomedical Sciences, and NIH National Institute of Allergy and Infectious Diseases T32AI132165.

## STAR Methods

### Resource Availability

#### Lead Contact

Further information and requests for resources and reagents should be directed to and will be fulfilled by the lead contact, George Vasmatzis.

#### Materials Availability

This study did not generate new unique reagents.

#### Data and Code Availability

- Raw data used in this study cannot be shared to preserve patient confidentiality, but derived data reported in this paper will be shared by the lead contact upon request
- This paper does not report original code
- Any additional information required to reanalyze the data reported in this paper is available from the lead contact upon request

### Experimental Model and Study Participant Details

All methods were carried out in accordance with relevant guidelines and regulations and all experimental protocols were approved by the Mayo Clinic institutional review board (IRB) studies: 15-005545 or 19-005326. Informed consent was obtained from all the participants and/ or their legal guardians. All data for this study was generated from tissue collected from female patients due to the focus on endometrial cancer.

### Method Details

#### MPseq bioinformatics analysis

Our bioinformatics pipeline reported structural variants from Mate-Pair library preparations (MPseq), a low-pass whole-genome DNA sequencing method as previously reported ^(8–11)^. Briefly, the sequenced fragments were mapped to reference genome GRCh38 by BIMA ^(50)^ and assessed for junctions and copy number variants by SVAtools. Junctions were detected from clusters of discordant (based on the mapped location and/or orientation of the reads in each fragment) fragments spanning two breakpoints ^(8)^. Copy number variants were called based on read-depth of concordant fragments ^(9)^. Junctions and CNVs were graphically illustrated for further inspection using genome, junction and region plots.

#### Detection and distinction of tandem and non-tandem duplications

Previously in WGS analysis of TDP in several cancers, including HCN endometrial cancer, the specific orientation of structural variants were employed to separate tandem duplications (intra-chromosomal 3-to-5 orientation) from the deletions, inversion, and inter-chromosomal translocations. However, upon analysis of the data in our cohort using whole genome visualization we were able to determine the widespread occurrence of duplications including 2+ distant intra- and inter-chromosomal locations. The typical tandem duplication analysis would characterize these primarily as simple translocations and would not include them in the final analysis. For this reason, we opted for a hybrid CNV-SV approach.

First, we treated each sample like an array and performed focal (<3.4Mb) duplication analysis. We looked for regions of copy number change that were smaller than a reasonable upper limit consistent with literature on TDP and had a higher copy number level than the region on either side, suggesting the possible presence of a focal gain. We then looked for the presence or absence of breakpoints on either side of the copy number change. We required that a valid duplication event fit the expected copy number change criteria, while also having at least one side correspond to the location of a breakpoint. This eliminated dubious copy number regions that were likely the results of polymorphic regions of the genome and would have no detectable breakpoint junctions explaining the focal CNV change, while including duplications that arose from other mechanisms other than a strict tandem duplication orientation.

This strategy increased our detection of duplication regions across the HCN endometrial cancer samples. This suggests that the TDP includes not just tandem duplications, but also regions of duplication that occur because of complex intra- and inter-chromosomal duplications. Most importantly this showed a distinct increase in the number of duplications that altered important cancer genes. We found that while it might be tempting to employ the specificity of the structural variant detection in WGS to strictly limit the allowed regions of gain in a duplication analysis, this type of analysis is at risk of missing important duplication regions. We found that the hybrid CNV-SV approach, a process similar to the copy number variant detection used to analyze array results supplemented by the SV breakpoints from the WGS analysis, was more accurate.

#### Cell isolation and drug response testing

Human endometrial tumor tissue was dissociated into single-cell suspensions using the Tumor Dissociation Kit, human (Miltenyi Biotec, 130-095-929) and the gentleMACS™ Dissociator (Miltenyi Biotech, 130-093-235) according to the manufacturer’s instructions. An equal number of cells (5x103) were seeded at 40 μl per well of a 96-well AkuraTM PLUS hanging drop plate (Insphero, CS-06-004-02) and incubated at 37°C for 6 days in DMEM/F-12 media containing 10% heat-inactivated horse serum, 5μg/ml insulin, 10ng/ml EGF, 10 μM Y-27632, and pen/strep allowing for the formation of microcancers. Microcancers from each well were then transferred to the corresponding well of a Corning® 96-well Clear Round Bottom Ultra-Low Attachment Microplate (Corning, 7007) and exposed to afatinib in varying concentrations and to the DMSO vehicle control for 6 days. Cell viability was measured using the CellTiter-Glo® assay (Promega, G7570) and expressed as ATP levels which are interpolated from an ATP standard curve.

#### Quantification and Statistical Analysis

Statistical analysis of tier distributions and drug response data were performed using GraphPad Prism v9.0.0. For drug response, interpolated ATP levels were normalized to DMSO control (= 100% viability), and drug dose-response curves were visualized using a nonlinear regression method (log(inhibitor) vs. normalized response – variable slope). The absolute half maximal inhibitory concentration (Abs IC50) values were interpolated. For tier distributions, Chi-square and Shapiro-Wilk testing were used to assess distribution and normality. See figure legends for details.

## Key Resources

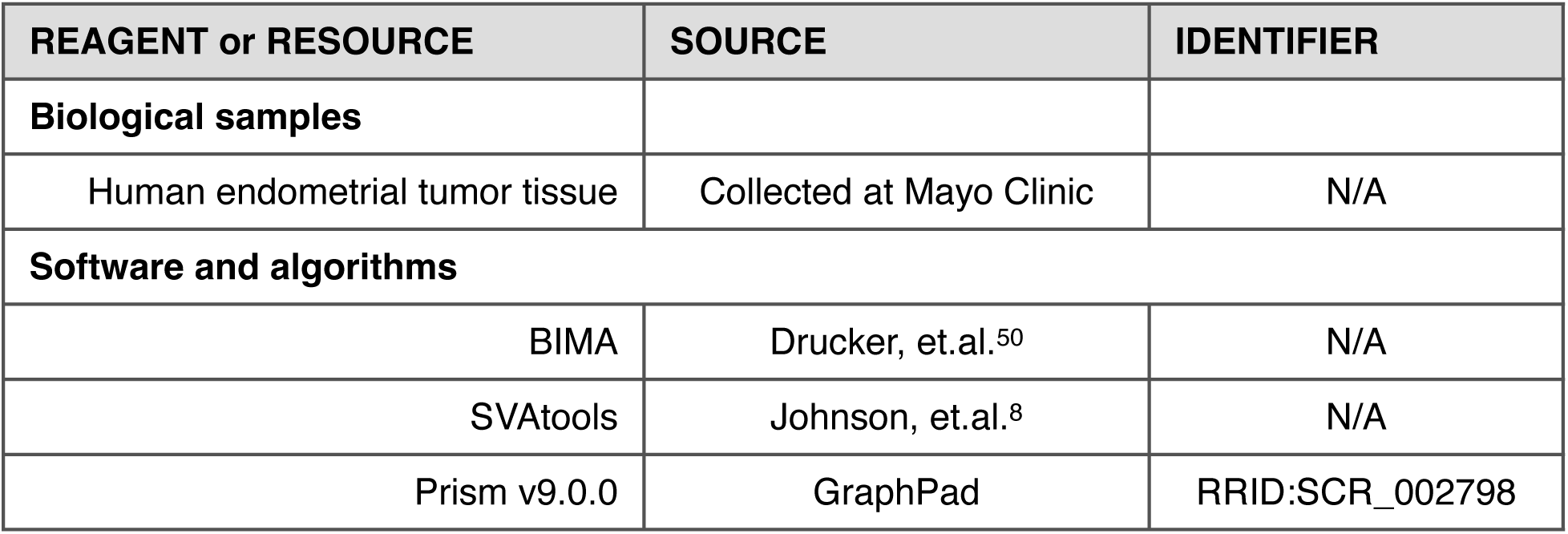

| REAGENT or RESOURCE | SOURCE | IDENTIFIER |
| --- | --- | --- |
| **Biological samples** |  |  |
| Human endometrial tumor tissue | Collected at Mayo Clinic | N/A |
| **Software and algorithms** |  |  |
| BIMA | Drucker, et.al.<sup>50</sup> | N/A |
| SVAtools | Johnson, et.al.<sup>8</sup> | N/A |
| Prism v9.0.0 | GraphPad | RRID:SCR_002798 |

## References

1 Bokhman, J.V. (1983). Two pathogenetic types of endometrial carcinoma. Gynecol Oncol 15, 10–17. 10.1016/0090-8258(83)90111-7.

2 Lobo, F.D., and Thomas, E. (2016). Type II endometrial cancers: A case series. J Midlife Health 7, 69–72. 10.4103/0976-7800.185335.

3 Kandoth, C., McLellan, M.D., Vandin, F., Ye, K., Niu, B., Lu, C., Xie, M., Zhang, Q., McMichael, J.F., Wyczalkowski, M.A., et al. (2013). Mutational landscape and significance across 12 major cancer types. Nature 502, 333–339. 10.1038/nature12634.

4 Menghi, F., Inaki, K., Woo, X., Kumar, P.A., Grzeda, K.R., Malhotra, A., Yadav, V., Kim, H., Marquez, E.J., Ucar, D., et al. (2016). The tandem duplicator phenotype as a distinct genomic configuration in cancer. Proc Natl Acad Sci U S A 113, E2373–2382. 10.1073/pnas.1520010113.

5 Xing, R., Zhou, Y., Yu, J., Yu, Y., Nie, Y., Luo, W., Yang, C., Xiong, T., Wu, W.K.K., Li, Z., et al. (2019). Whole-genome sequencing reveals novel tandem-duplication hotspots and a prognostic mutational signature in gastric cancer. Nat Commun 10, 2037. 10.1038/s41467-019-09644-6.

6 Rao, M., and Powers, S. (2018). Tandem Duplications May Supply the Missing Genetic Alterations in Many Triple-Negative Breast and Gynecological Cancers. Cancer Cell 34, 179–180. 10.1016/j.ccell.2018.07.011.

7 Menghi, F., Barthel, F.P., Yadav, V., Tang, M., Ji, B., Tang, Z., Carter, G.W., Ruan, Y., Scully, R., Verhaak, R.G.W., et al. (2018). The Tandem Duplicator Phenotype Is a Prevalent Genome-Wide Cancer Configuration Driven by Distinct Gene Mutations. Cancer Cell 34, 197–210.e5. 10.1016/j.ccell.2018.06.008.

8 Johnson, S.H., Smadbeck, J.B., Smoley, S.A., Gaitatzes, A., Murphy, S.J., Harris, F.R., Drucker, T.M., Zenka, R.M., Pitel, B.A., Rowsey, R.A., et al. (2018). SVAtools for junction detection of genome-wide chromosomal rearrangements by mate-pair sequencing (MPseq). Cancer Genet 221, 1–18. 10.1016/j.cancergen.2017.11.009.

9 Smadbeck, J.B., Johnson, S.H., Smoley, S.A., Gaitatzes, A., Drucker, T.M., Zenka, R.M., Kosari, F., Murphy, S.J., Hoppman, N., Aypar, U., et al. (2018). Copy number variant analysis using genome-wide mate-pair sequencing. Genes Chromosomes Cancer 57, 459–470. 10.1002/gcc.5.

10 Mansfield, A.S., Murphy, S.J., Harris, F.R., Robinson, S.I., Marks, R.S., Johnson, S.H., Smadbeck, J.B., Halling, G.C., Yi, E.S., Wigle, D., et al. (2016). Chromoplectic TPM3-ALK rearrangement in a patient with inflammatory myofibroblastic tumor who responded to ceritinib after progression on crizotinib. Ann Oncol 27, 2111–2117. 10.1093/annonc/mdw405.

11 Murphy, S.J., Hart, S.N., Halling, G.C., Johnson, S.H., Smadbeck, J.B., Drucker, T., Lima, J.F., Rohakhtar, F.R., Harris, F.R., Kosari, F., et al. (2016). Integrated Genomic Analysis of Pancreatic Ductal Adenocarcinomas Reveals Genomic Rearrangement Events as Significant Drivers of Disease. Cancer Res 76, 749–761. 10.1158/0008-5472.CAN-15-2198.

12 Carvajal-Garcia, J., Cho, J.-E., Carvajal-Garcia, P., Feng, W., Wood, R.D., Sekelsky, J., Gupta, G.P., Roberts, S.A., and Ramsden, D.A. (2020). Mechanistic basis for microhomology identification and genome scarring by polymerase theta. Proc Natl Acad Sci U S A 117, 8476–8485. 10.1073/pnas.1921791117.

13 Moore, G., Majumdar, R., Powell, S.N., Khan, A.J., Weinhold, N., Yin, S., and Higginson, D.S. (2022). Templated Insertions Are Associated Specifically with BRCA2 Deficiency and Overall Survival in Advanced Ovarian Cancer. Mol Cancer Res 20, 1061–1070. 10.1158/1541-7786.MCR-21-1012.

14 Chakravarty, D., Gao, J., Phillips, S., Kundra, R., Zhang, H., Wang, J., Rudolph, J.E., Yaeger, R., Soumerai, T., Nissan, M.H., et al. (2017). OncoKB: A Precision Oncology Knowledge Base. JCO Precision Oncology, 1–16. 10.1200/PO.17.00011.

15 Tate, J.G., Bamford, S., Jubb, H.C., Sondka, Z., Beare, D.M., Bindal, N., Boutselakis, H., Cole, C.G., Creatore, C., Dawson, E., et al. (2019). COSMIC: the Catalogue Of Somatic Mutations In Cancer. Nucleic Acids Research 47, D941–D947. 10.1093/nar/gky1015.

16 Piñero, J., Ramírez-Anguita, J.M., Saüch-Pitarch, J., Ronzano, F., Centeno, E., Sanz, F., and Furlong, L.I. (2019). The DisGeNET knowledge platform for disease genomics: 2019 update. Nucleic Acids Research, gkz1021. 10.1093/nar/gkz1021.

17 Becker, K.G., Barnes, K.C., Bright, T.J., and Wang, S.A. (2004). The genetic association database. Nat Genet 36, 431–432. 10.1038/ng0504-431.

18 Kelman, G., Brandes, N., and Linial, M. (2021). The FABRIC Cancer Portal: A Ranked Catalogue of Gene Selection in Tumors Over the Human Coding Genome. Cancer Research 81, 1178–1185. 10.1158/0008-5472.CAN-20-3147.

19 Huret, J.-L., Ahmad, M., Arsaban, M., Bernheim, A., Cigna, J., Desangles, F., Guignard, J.- C., Jacquemot-Perbal, M.-C., Labarussias, M., Leberre, V., et al. (2013). Atlas of genetics and cytogenetics in oncology and haematology in 2013. Nucleic Acids Res 41, D920–924. 10.1093/nar/gks1082.

20 Vasmatzis, G., Liu, M.C., Reganti, S., Feathers, R.W., Smadbeck, J., Johnson, S.H., Schaefer Klein, J.L., Harris, F.R., Yang, L., Kosari, F., et al. (2020). Integration of Comprehensive Genomic Analysis and Functional Screening of Affected Molecular Pathways to Inform Cancer Therapy. Mayo Clinic Proceedings 95, 306–318. 10.1016/j.mayocp.2019.07.019.

21 Ganguli, A., Mostafa, A., Saavedra, C., Kim, Y., Le, P., Faramarzi, V., Feathers, R.W., Berger, J., Ramos-Cruz, K.P., Adeniba, O., et al. (2021). Three-dimensional microscale hanging drop arrays with geometric control for drug screening and live tissue imaging. Sci Adv 7, eabc1323. 10.1126/sciadv.abc1323.

22 Bonazzoli, E., Cocco, E., Lopez, S., Bellone, S., Zammataro, L., Bianchi, A., Manzano, A., Yadav, G., Manara, P., Perrone, E., et al. (2019). PI3K oncogenic mutations mediate resistance to afatinib in HER2/neu overexpressing gynecological cancers. Gynecol Oncol 153, 158–164. 10.1016/j.ygyno.2019.01.002.

23 Zhou, L., Ren, Y., Wang, X., Miao, D., Lizaso, A., Li, H., Han-Zhang, H., Qian, J., and Yang, H. (2019). Efficacy of afatinib in a HER2 amplification-positive endometrioid adenocarcinoma patient- a case report. Onco Targets Ther 12, 5305–5309. 10.2147/OTT.S206732.

24 Li, C.G., Nyman, J.E., Braithwaite, A.W., and Eccles, M.R. (2011). PAX8 promotes tumor cell growth by transcriptionally regulating E2F1 and stabilizing RB protein. Oncogene 30, 4824–4834. 10.1038/onc.2011.190.

25 Bleu, M., Mermet-Meillon, F., Apfel, V., Barys, L., Holzer, L., Bachmann Salvy, M., Lopes, R., Amorim Monteiro Barbosa, I., Delmas, C., Hinniger, A., et al. (2021). PAX8 and MECOM are interaction partners driving ovarian cancer. Nat Commun 12, 2442. 10.1038/s41467-021-22708-w.

26 Jongen, V., Briët, J., de Jong, R., ten Hoor, K., Boezen, M., van der Zee, A., Nijman, H., and Hollema, H. (2009). Expression of estrogen receptor-alpha and -beta and progesterone receptor-A and -B in a large cohort of patients with endometrioid endometrial cancer. Gynecologic Oncology 112, 537–542. 10.1016/j.ygyno.2008.10.032.

27 Shen, F., Gao, Y., Ding, J., and Chen, Q. (2017). Is the positivity of estrogen receptor or progesterone receptor different between type 1 and type 2 endometrial cancer? Oncotarget 8, 506–511. 10.18632/oncotarget.13471.

28 Wang, Y.-P., Guo, P.-T., Zhu, Z., Zhang, H., Xu, Y., Chen, Y.-Z., Liu, F., and Ma, S.-P. (2017). Pleomorphic adenoma gene like-2 induces epithelial-mesenchymal transition via Wnt/β- catenin signaling pathway in human colorectal adenocarcinoma. Oncol Rep 37, 1961–1970. 10.3892/or.2017.5485.

29 Wu, L., Zhou, Z., Han, S., Chen, J., Liu, Z., Zhang, X., Yuan, W., Ji, J., and Shu, X. (2020). PLAGL2 promotes epithelial-mesenchymal transition and mediates colorectal cancer metastasis via β-catenin-dependent regulation of ZEB1. Br J Cancer 122, 578–589. 10.1038/s41416-019-0679-z.

30 Hubalek, M.M., Widschwendter, A., Erdel, M., Gschwendtner, A., Fiegl, H.M., Müller, H.M., Goebel, G., Mueller-Holzner, E., Marth, C., Spruck, C.H., et al. (2004). Cyclin E dysregulation and chromosomal instability in endometrial cancer. Oncogene 23, 4187–4192. 10.1038/sj.onc.1207560.

31 Nakayama, K., Rahman, M.T., Rahman, M., Nakamura, K., Ishikawa, M., Katagiri, H., Sato, E., Ishibashi, T., Iida, K., Ishikawa, N., et al. (2016). CCNE1 amplification is associated with aggressive potential in endometrioid endometrial carcinomas. International Journal of Oncology 48, 506–516. 10.3892/ijo.2015.3268.

32 Chan, A.M., Enwere, E., McIntyre, J.B., Wilson, H., Nwaroh, C., Wiebe, N., Ou, Y., Liu, S., Wiedemeyer, K., Rambau, P.F., et al. (2020). Combined CCNE1 high-level amplification and overexpression is associated with unfavourable outcome in tubo-ovarian high-grade serous carcinoma. J Pathol Clin Res 6, 252–262. 10.1002/cjp2.168.

33 Cherniack, A.D., Shen, H., Walter, V., Stewart, C., Murray, B.A., Bowlby, R., Hu, X., Ling, S., Soslow, R.A., Broaddus, R.R., et al. (2017). Integrated Molecular Characterization of Uterine Carcinosarcoma. Cancer Cell 31, 411–423. 10.1016/j.ccell.2017.02.010.

34 Zhao, S., Choi, M., Overton, J.D., Bellone, S., Roque, D.M., Cocco, E., Guzzo, F., English, D.P., Varughese, J., Gasparrini, S., et al. (2013). Landscape of somatic single-nucleotide and copy-number mutations in uterine serous carcinoma. Proc. Natl. Acad. Sci. U.S.A. 110, 2916–2921. 10.1073/pnas.1222577110.

35 Xu, H., George, E., Kinose, Y., Kim, H., Shah, J.B., Peake, J.D., Ferman, B., Medvedev, S., Murtha, T., Barger, C.J., et al. (2021). CCNE1 copy number is a biomarker for response to combination WEE1-ATR inhibition in ovarian and endometrial cancer models. Cell Reports Medicine 2, 100394. 10.1016/j.xcrm.2021.100394.

36 Zheng, X., Chen, L., Liu, W., Zhao, S., Yan, Y., Zhao, J., Tian, W., and Wang, Y. (2023). CCNE1 is a predictive and immunotherapeutic indicator in various cancers including UCEC: a pan-cancer analysis. Hereditas 160, 13. 10.1186/s41065-023-00273-0.

37 Li, J., Bennett, K., Stukalov, A., Fang, B., Zhang, G., Yoshida, T., Okamoto, I., Kim, J.-Y., Song, L., Bai, Y., et al. (2013). Perturbation of the mutated EGFR interactome identifies vulnerabilities and resistance mechanisms. Mol Syst Biol 9, 705. 10.1038/msb.2013.61.

38 Janes, P.W., Lackmann, M., Church, W.B., Sanderson, G.M., Sutherland, R.L., and Daly, R.J. (1997). Structural determinants of the interaction between the erbB2 receptor and the Src homology 2 domain of Grb7. J Biol Chem 272, 8490–8497. 10.1074/jbc.272.13.8490.

39 Fiddes, R.J., Campbell, D.H., Janes, P.W., Sivertsen, S.P., Sasaki, H., Wallasch, C., and Daly, R.J. (1998). Analysis of Grb7 recruitment by heregulin-activated erbB receptors reveals a novel target selectivity for erbB3. J Biol Chem 273, 7717–7724. 10.1074/jbc.273.13.7717.

40 Liu, C.-X., Qiao, X.-J., Xing, Z.-W., and Hou, M.-X. (2018). The SNORA21 expression is upregulated and acts as a novel independent indicator in human gastric cancer prognosis. European Review for Medical and Pharmacological Sciences 22, 5519–5524. 10.26355/eurrev_201809_15812.

41 Yoshida, K., Toden, S., Weng, W., Shigeyasu, K., Miyoshi, J., Turner, J., Nagasaka, T., Ma, Y., Takayama, T., Fujiwara, T., et al. (2017). SNORA21 – An Oncogenic Small Nucleolar RNA, with a Prognostic Biomarker Potential in Human Colorectal Cancer. EBioMedicine 22, 68–77. 10.1016/j.ebiom.2017.07.009.

42 Kim, D.-S., Camacho, C.V., Nagari, A., Malladi, V.S., Challa, S., and Kraus, W.L. (2019). Activation of PARP-1 by snoRNAs Controls Ribosome Biogenesis and Cell Growth via the RNA Helicase DDX21. Molecular Cell 75, 1270–1285.e14. 10.1016/j.molcel.2019.06.020.

43 Li, W., Li, H., Zhang, L., Hu, M., Li, F., Deng, J., An, M., Wu, S., Ma, R., Lu, J., et al. (2017). Long non-coding RNA LINC00672 contributes to p53 protein-mediated gene suppression and promotes endometrial cancer chemosensitivity. J Biol Chem 292, 5801–5813. 10.1074/jbc.M116.758508.

44 Tang, J., Zhuo, H., Zhang, X., Jiang, R., Ji, J., Deng, L., Qian, X., Zhang, F., and Sun, B. (2014). A novel biomarker Linc00974 interacting with KRT19 promotes proliferation and metastasis in hepatocellular carcinoma. Cell Death Dis 5, e1549. 10.1038/cddis.2014.518.

45 Su, H., Xu, T., Ganapathy, S., Shadfan, M., Long, M., Huang, T.H.-M., Thompson, I., and Yuan, Z.-M. (2014). Elevated snoRNA biogenesis is essential in breast cancer. Oncogene 33, 1348–1358. 10.1038/onc.2013.89.

46 Blenkiron, C., Hurley, D.G., Fitzgerald, S., Print, C.G., and Lasham, A. (2013). Links between the Oncoprotein YB-1 and Small Non-Coding RNAs in Breast Cancer. PLoS ONE 8, e80171. 10.1371/journal.pone.0080171.

47 Mourksi, N.-E.-H., Morin, C., Fenouil, T., Diaz, J.-J., and Marcel, V. (2020). snoRNAs Offer Novel Insight and Promising Perspectives for Lung Cancer Understanding and Management. Cells 9, 541. 10.3390/cells9030541.

48 Tseng, Y.-Y., Moriarity, B.S., Gong, W., Akiyama, R., Tiwari, A., Kawakami, H., Ronning, P., Reuland, B., Guenther, K., Beadnell, T.C., et al. (2014). PVT1 dependence in cancer with MYC copy-number increase. Nature 512, 82–86. 10.1038/nature13311.

49 Khurana, E., Fu, Y., Chakravarty, D., Demichelis, F., Rubin, M.A., and Gerstein, M. (2016). Role of non-coding sequence variants in cancer. Nat Rev Genet 17, 93–108. 10.1038/nrg.2015.17.

50 Drucker, T.M., Johnson, S.H., Murphy, S.J., Cradic, K.W., Therneau, T.M., and Vasmatzis, G. (2014). BIMA V3: an aligner customized for mate pair library sequencing. Bioinformatics 30, 1627–1629. 10.1093/bioinformatics/btu078.

