## Supplemental Figures for "Functional impact of the hyperduplication genomophenotype in high copy number endometrial cancer"

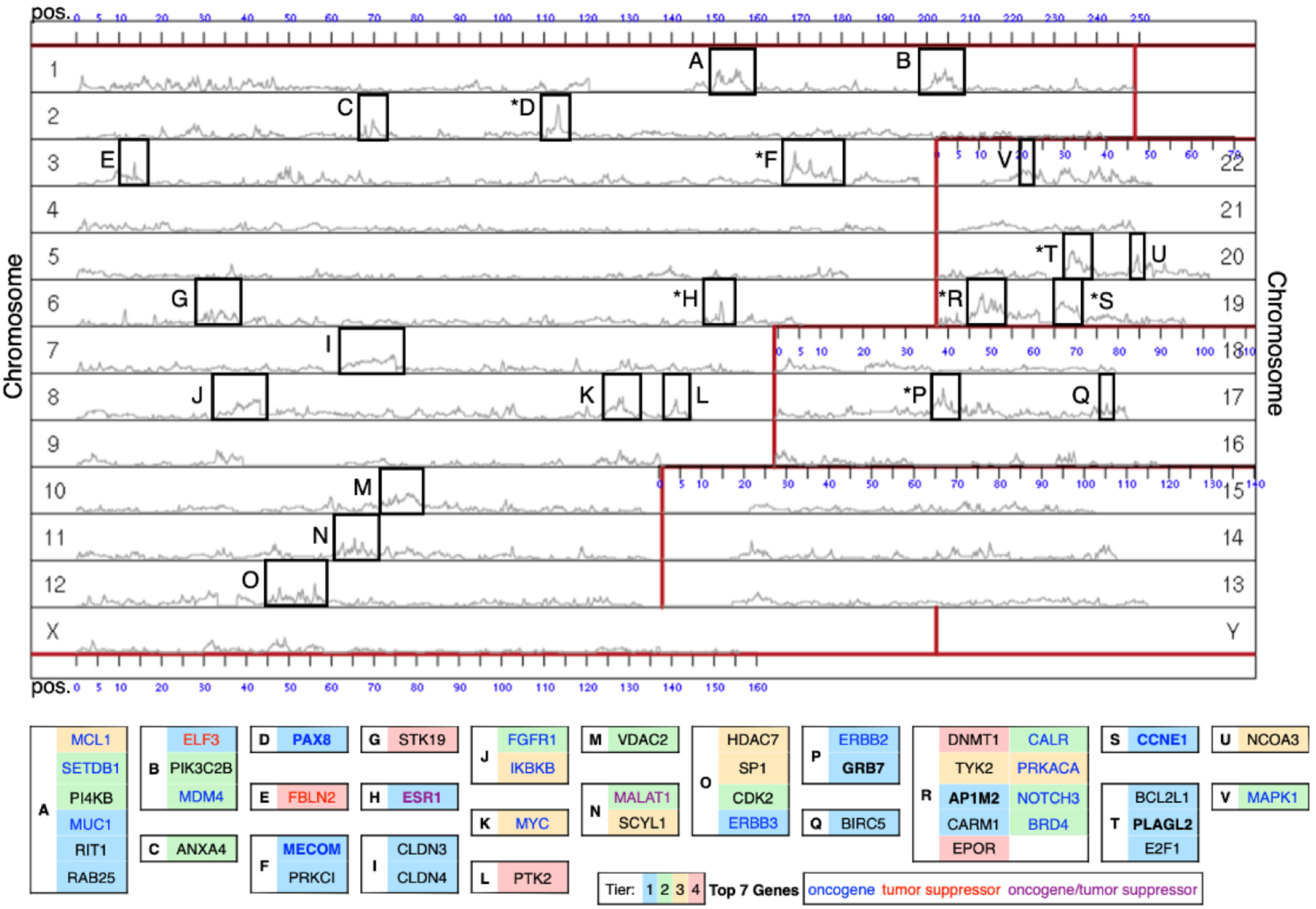


**Supplemental Figure S1. A genome U-plot showing regional frequency of duplications.** Chromosomes are arranged in a U shape and numbered along the left and right sides; the scales at the top and bottom indicate position. The gray line indicates the frequency of duplication across our HDGP-EC data set. Letters indicate high-frequency RDMRs and correspond to the table, which highlights genes of interest in each region. Gene boxes are colored by Tier, and Top 7 Genes are in bold. Blue text denotes an oncogene, red text denotes a tumor suppressor, and purple text denotes an oncogene/tumor suppressor as determined by the COSMIC Cancer Gene Census. *Further detailed in Supplemental Figure S3.

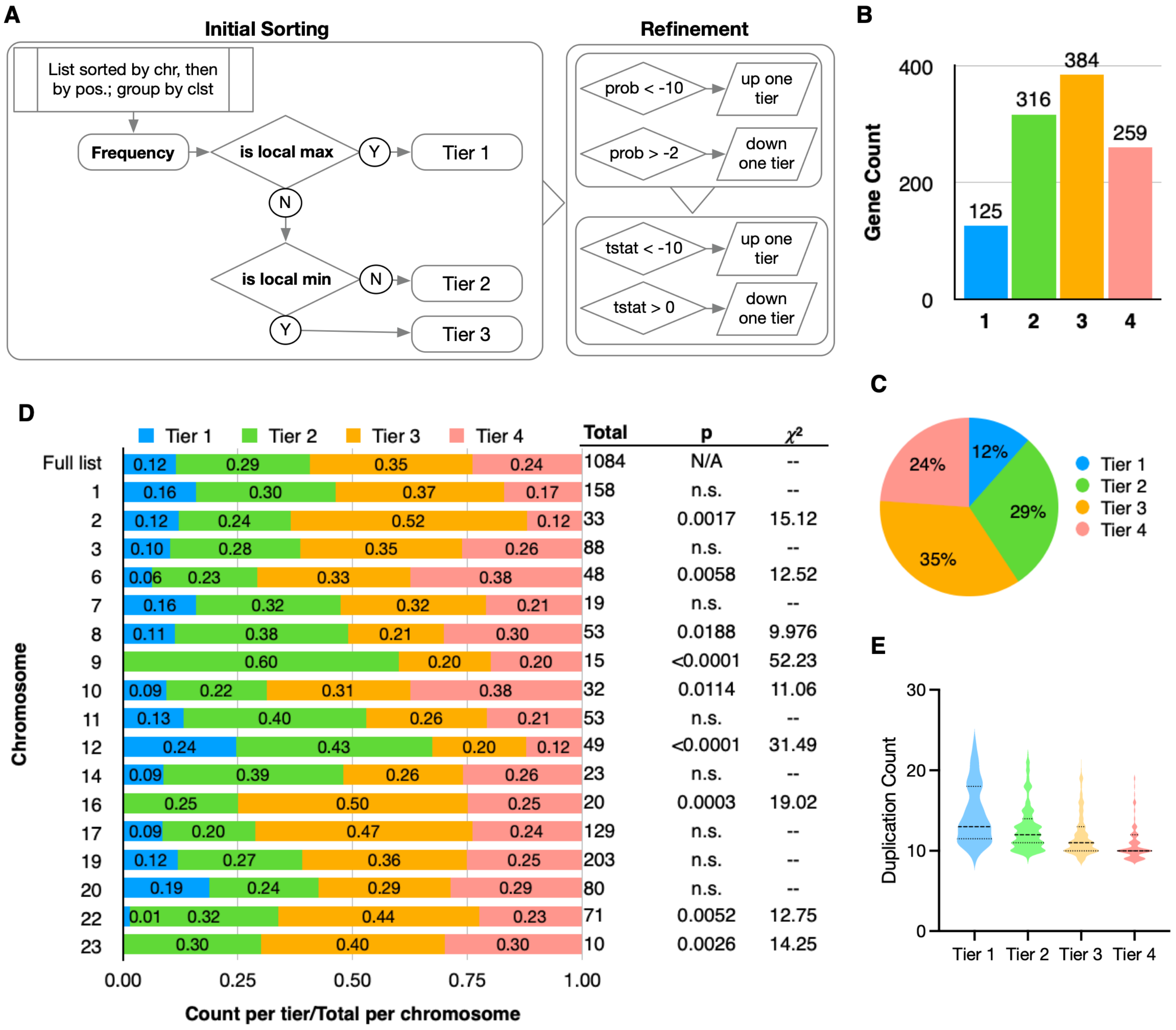


**Supplemental Figure S2. A tier system was developed to allow deeper analysis through hierarchical grouping. A)** Preliminary sorting was based on each gene's frequency within the HCNEC data set in the context of its genomic region (pos indicates position, clst indicates cluster). Tiers were refined based on probability (prob) and t-statistic (tstat) related to expression data versus non-EC cases in the larger overall data set. **B,C)** The resulting tiers vary in size and reflect a normal distribution. Shapiro-Wilk test, W = 0.9695, p = 0.8386, alpha = 0.05. **D)** The distribution of tiers across several chromosomes differs significantly from that of the full list of hyper-duplicated genes. Chromosome 12 is overrepresented in Tier 1 and Tier 2; the others are overrepresented in Tiers 3 and 4. Chi-square test, alpha = 0.05, DF = 3. The frequency of duplications on chromosomes 4, 5, 13, 15, 18, and 21 fell below the significance cutoff of 9 for inclusion in the list. **E)** A violin plot based on the duplication count in each tier shows the effect of expression-based refinement and reveals a bimodal distribution in Tier 1. The dashed line indicates median, the dotted lines indicate quartiles.

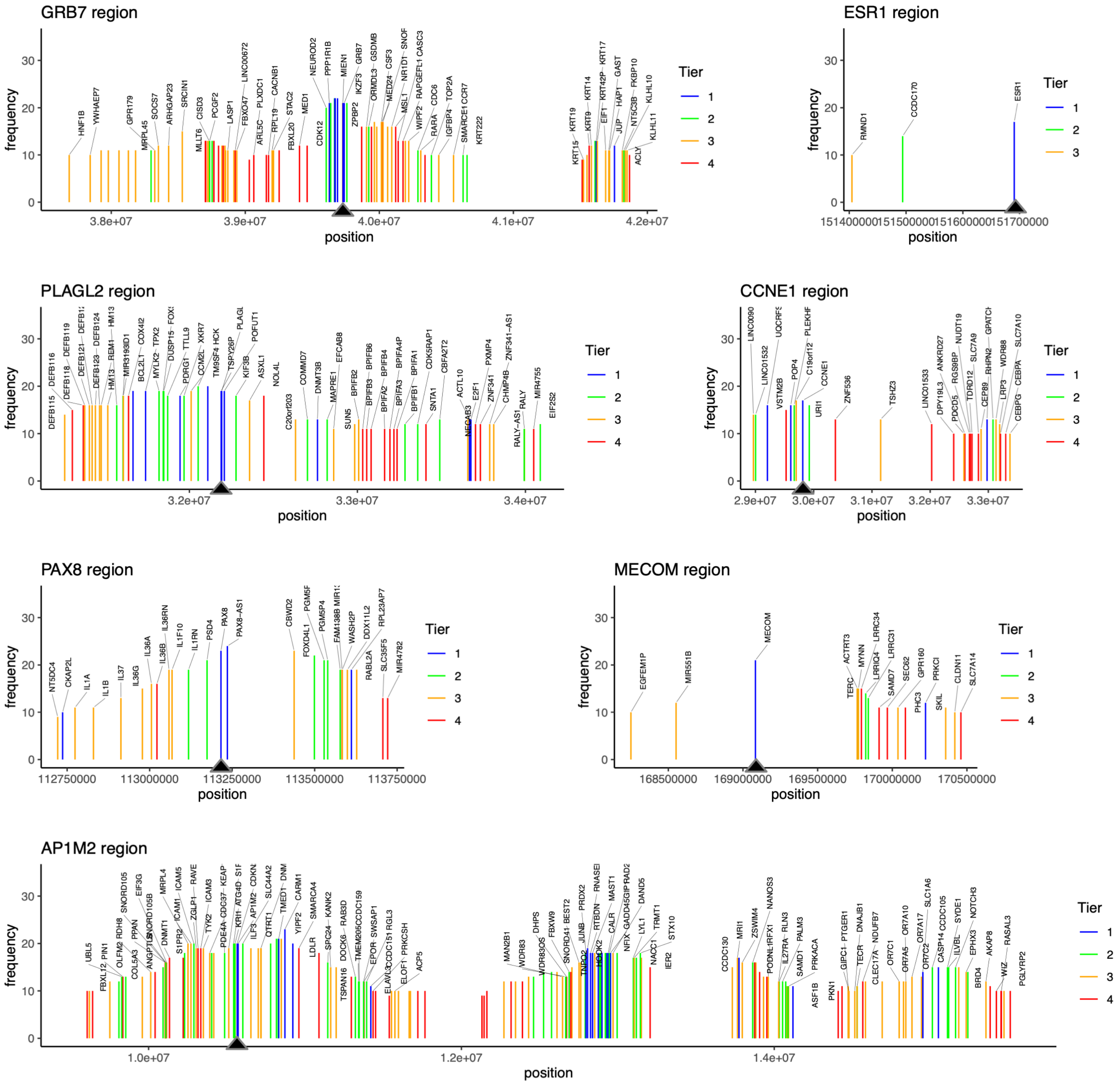


**Supplemental Figure S3. The Top 7 Genes in their regional context, colored by Tier.** Small arrows indicate the gene of interest. While the duplicated genes vary in size and density, most contain other genes with either known cancer associations or known functions in cancer hallmark pathways. *ESR1* is the only top gene to appear as the only peak in a hyperduplicated region. Some labels were omitted to preserve readability.
